# A novel virtual plus-maze for studying electrophysiological correlates of spatial reorientation

**DOI:** 10.1101/369207

**Authors:** Ágoston Török, Andrea Kóbor, Ferenc Honbolygó, Travis Baker

## Abstract

Quick reorientation is an essential part of successful navigation. Despite growing attention to this ability, little is known about how reorientation happens in humans. To this aim, we recorded EEG from 34 participants. Participants were navigating a simple virtual reality plus-maze where at the beginning of each trial they were randomly teleported to either the North or the South alley. Results show that the teleportation event caused a quick reorientation effect over occipito-parietal areas as early as 100 msec; meaning that despite the known stochastic nature of the teleportation, participants built up expectations for their place of arrival. This result has important consequences for the optimal design of Virtual reality locomotion.

## 1. Introduction

Regaining our spatial orientation in an environment (e.g. while walking distracted by our cell phone) is essential for everyday navigation. While the neural mechanisms underlying spatial orientation are well described in animal models, compelling evidence linking neural activity with reorientation in humans remains elusive. In order to examine the effect of reorientation in humans, we recorded the encephalogram (EEG) while teleporting human subjects to different starting locations in a virtual plus-maze, the gold standard to assess reorientation behavior in rodents Packard & McGaugh (1996).

In this experiment, participants’ start position was first consistent across trials (always starting from the South alley) and then in a test phase became inconsistent across trials (random teleportation events to North or South starting alleys). Given that the navigation system of rodents is highly sensitive to such reorientation effects in the plus-maze, we expected that the reorientation in the virtual plus-maze may have similar effects and may therefore be visible in the EEG. Given the novelty of this approach, we refrain from making any specific prediction. Nevertheless, these results here provide insights into human navigation processes related to reorientation during navigation and illustrate an approach for future research on combing EEG with virtual navigation as a means to parallel animal and computational models of spatial navigation.

## 2. Materials and methods

### 2.1 Participants

Thirty-four participants (16 males; 34 right-handed; aged 19–29, *M* = 22, *SD* = 2.6) with normal or corrected-to-normal vision participated in the experiment. Participants were undergraduate students recruited from the Budapest University of Technology and Economics and the Eötvös Loránd University and each received course credit or a monetary compensation for their participation. All participants gave informed consent. The study was approved by the local research ethics committee and was conducted in accordance with the ethical standards prescribed in the 1964 Declaration of Helsinki.

### 2.2 Procedure

Participants started either in point *α* or *β* (see Fig 1) and were asked to choose between the two horizontal alleys. Following their choice, they were translated and rotated to the chosen alley, followed by the feedback stimulus. The details of the reward collection can be found in the in our earlier study which contains the analysis of the reward phase Török et al. (2017).

**Figure 1:**
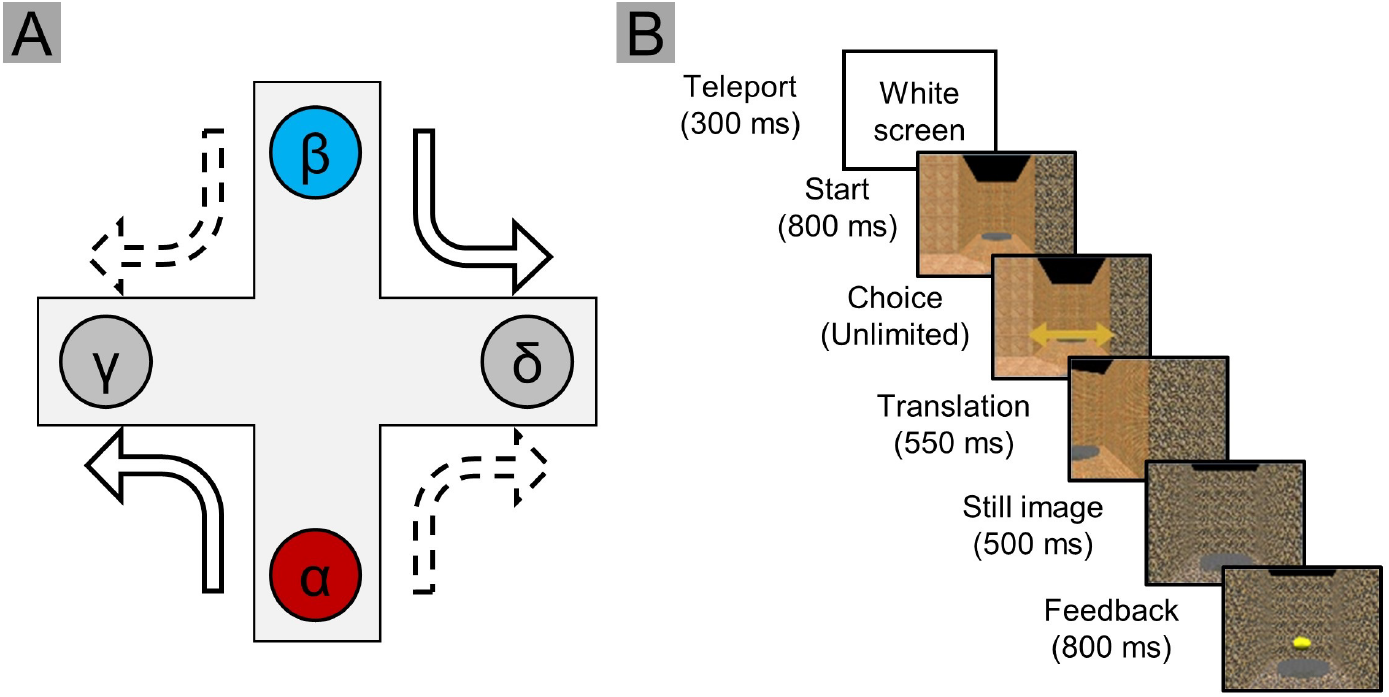
The layout of the plus-maze and the trial timeline. (A.) Participants started each trial randomly in either the *α* or *β* alley. (B.) Then they chose between the other two alleys, translated there and received a feedback (positive or neutral).

In order to establish a stable allocentric frame before the experiment Török et al. (2017), a practice phase of 130 trials was run where participants always started from the lower alley. After the practice phase, 4 blocks of 100 trials were recorded (total of 400 trials). The experiment lasted cca. 90 minutes with the electrode cap setup and debriefing. During the task, EEG was recorded from 62 sites placed according to the 10/20 system. The recording was done with BrainAmp amplifiers and MOVE system (Brain Products GmbH) with 1000 Hz sampling rate. An online 0.1-70 Hz bandpass filter was applied during acquisition. During recording all impedances were kept below 30 kΩ.

### 2.3 Data preprocessing

Preanalysis of the electrophysiological data was done using Matlab and EEGLAB Delorme & Makeig (2004). First, data were re-referenced to average reference Bertrand et al. (1985), and the original reference was retained (FCz). Then, we filtered the data with a 0.2-30 Hz band-pass FIR filter, epoched using a −100 msec and +500 msec window relative to the appearance of trial starts. Independent component analysis Delorme & Makeig (2004) and amplitude thresholding were used to reduce eye blinks and muscle artifacts. Detailed description of these steps can be found in Török et al. (2017).

### 2.4 Modeling and statistical analysis

#### 2.4.1 Behavioral data analysis

We calculated the median decision times in the task in the two possible starting positions and compared them in a Bayesian paired-samples *t*-test using JASP Team (2018). Median decision times were used since distribution was very much left skewed (Fig S1). The null hypothesis (*H*0) was that there is no difference in the decision times from the two starting positions, whereas the alternative hypothesis (*H*+) was that decisions take more time from the non-default starting alley. If we found evidence for the null, it would suggest smaller cognitive effort of reorientation, whereas if we found evidence for the alternative hypothesis, it would suggest larger cognitive effort. Following the objective Bayesian analysis routine Berger (2006), we specified 0.707 as the width of the half-Cauchy distribution prior. According to Wagenmakers et al. Wagenmakers et al. (2011), *BF_Alternative−Null_* values between 1 and 3 indicate anecdotal evidence for *H_Alternative_*, while values between 3 and 10 indicate substantial evidence for *H_Alternative_*.

#### 2.4.2 Electrophyisological data analysis

Because each trial started with the participant randomly placed in either the South or North alley (see Fig. 1), they had to reorient themselves every time. Therefore, we looked at whether the ERPs time-locked to the start events differ for the two starting positions. As a data-driven approach, we calculated (1) global field power (GFP) and (2) topographic dissimilarity (TD) Lehmann & Skrandies (1980); Murray et al. (2008); Skrandies (1990) using RAGU (Randomization Graphical User interface; http://www.thomaskoenig.ch/Ragu.htm). Before the TD analysis, scalp topographies were normalized by the intensity of the signals at each time point; thus, significant results reflect pure topographic differences, probably driven by the involvement of new generators or change in the existing generators The threshold for all randomization based statistical testing was set to p < .05 based on 5000 iterations Koenig et al. (2011). The analysis was only performed for timepoints where the assumption of topographic consistency was not violated between subject according to TCT Koenig & Melie-García (2010). Furthermore, only differences meeting the Global Duration Statistics Criteria are reported.

After the topographic analysis, differences in topography were further explored on the electrodes where the scalp topography difference was greatest. Here, the results of the analyses are reported with False Discovery Rate (FDR) and Cluster method corrections applied Maris & Oostenveld (2007).

## 3 Results

### 3.1. Behavioral results

First, we investigated if the reorientation process had any behavioral correlates. Earlier analysis showed that participants were engaged in the task and were not choosing alleys randomly Török et al. (2017). Further, because during the practice phase participants were always starting in the South alley, they built up a strong a priori expectation for the starting alley Török et al. (2017). Consequently, they developed an intrinsic start facing preference in their internal map. We hypothesized that starting in the non-default (North) vs. in the default (South) would lead to longer decision times as a behavioral correlate of the reorientation process.

The analysis revealed moderate evidence for no difference in decision times (*BF*_+0_ = 0.302, error% < 0.001, *M_North_* = 753.0 msec, *SE_North_* = 52.32 msec, *M_South_* = 745.5 msec, *SE_South_* = 52.40 msec). The effect means that the hypothesis of no difference is approximately 3 times more likely based on the data than the alternative hypothesis of existing difference. As Figure 2 shows, evidence was favoring the null hypothesis all along the experiment.

**Figure 2:**
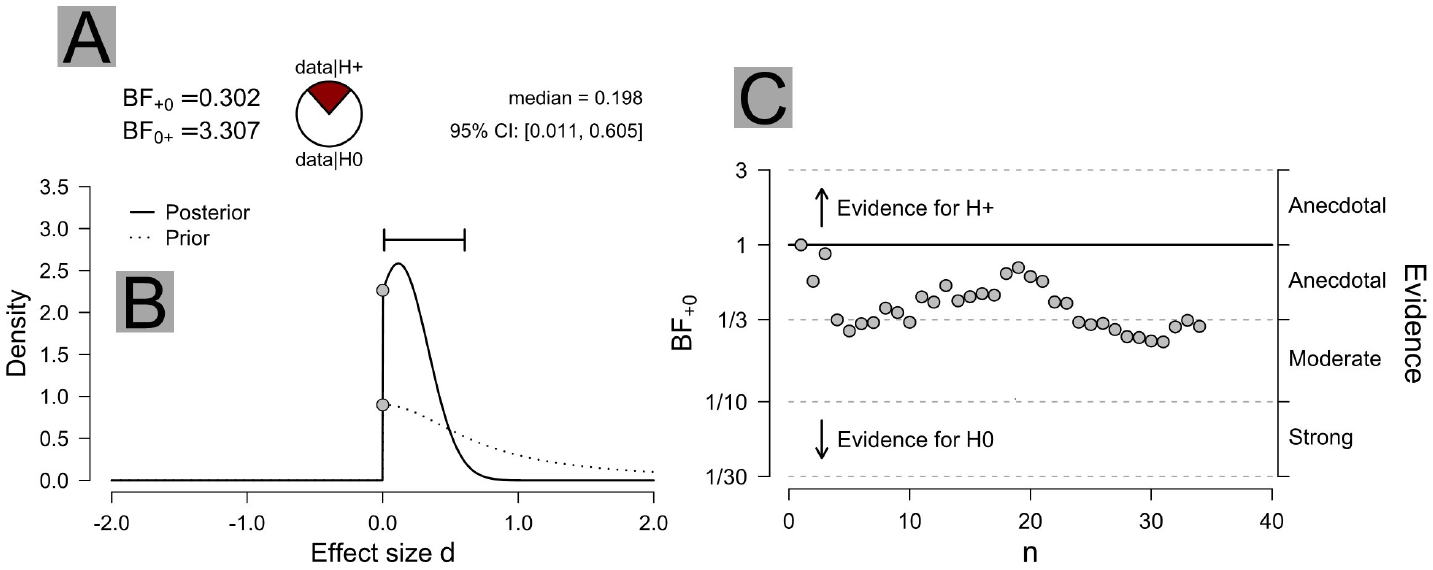
Results of the Bayesian t-test. A) The modeling found moderately strong evidence (*BF* ˜3) for the hypothesis that there is no decision times difference after starting in the unexpected vs expected alley. B) As can be seen, the posterior distribution largely overlaps with the prior distribution. C) sequential testing also shows that the results are more likely to support the null hypothesis with the addition of each participant.

Based on the behavioral data, we expected to find neural correlates of the reorientation process in earlier stages of processing that would not lead to increased decision time during the task.

### 3.2. Results of the topographic EEG analysis

#### 3.2.1. Results of the topographic dissimilarity analysis

Here, ERPs were compared between the expected vs. unexpected starting alley to find evidence for a reorientation process in the unexpected but not in the expected alley. First, differences in topographic dissimilarities were examined. The topographic dissimilarity analysis (TANOVA Murray et al. (2008)) revealed differences in scalp topographies between the two conditions. First, scalp topographies were significantly different between 103 and 134 msec (*p* < .05, see Fig. 3). The difference of the scalp topographies showed a large negativity over the parieto-occipital midline in the unexpected starting location Fig. 3). As the difference was maximal over the POz electrode, we analyzed waveforms here using both parametric testing (with FDR correction) and non-parametric testing (with Cluster method correction). Differences were found between 114 and 163 msec, where a negative deflection is visible on the waveforms when participants started in the North alley (see Fig. S2). Additional differences were found between 179 and 197 msec.

**Figure 3:**
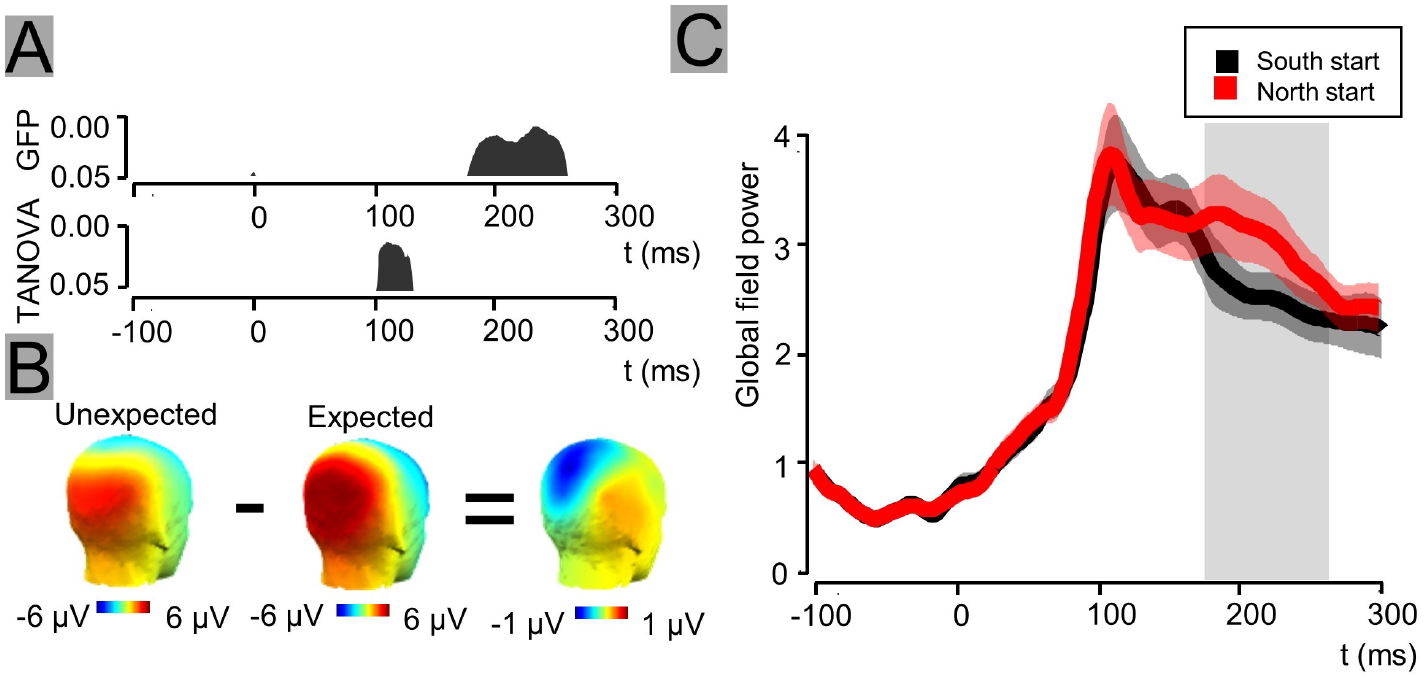
Reorientation at trial starts. A) Analysis revealed significant difference between scalp topographies from 103 to 134 msec and a global field power difference from 182 to 262 msec. These differences were significant after the application of Global Duration Statistics Criteria. B) The topographic difference is attributable to a negative deflection over parieto-occipital sites in the unexpected starting alley. C) The global field power difference means that there the same topographies are more pronounced in the time window of 182-262 msec

#### 3.2.2. Results of the global field power analysis

Next, we conducted analysis on the global field power values. Since topographic dissimilarity revealed an activity around 100 msec, we expected difference in the global field power in this or later time windows showing a cognitive effort to accommodate the unexpected North starting position. In line with our expectation, the analysis revealed difference in global field power from 181 to 264 msec (*p* < .05, see Fig. 3), where the scalp field power was stronger for North starting position.

In general, the difference in global field power means a stronger presence of the same scalp topography in one condition. Here it showed stronger activation in the right lateralized parieto-occipital processing with a peak over the O2 electrode (see Fig. S3). The difference was also the greatest over this electrode. Analysis of the waveforms was done using parametric testing (with FDR correction) and non-parametric testing (with Cluster method correction) on this electrode and showed significant differences between 123 msec and 152 msec and between 175 and 300 msec.

To rule out the more simplistic explanation of pure texture related differences without spatial processing we did a control analysis (see Fig S4), which ruled out this alternative hypothesis.

## 4. Discussion

We presented electrophysiological evidence for early, reorientation-related processes in an experiment involving virtual teleportation to random starting position in the beginning of trials. We found that starting in the unexpected alley did not result in longer decision times but was correlated with an early (˜100 msec) medial parieto-occipital negativity and a later increased bilateral occipital activity in the EEG waveforms. The present results are the first scalp EEG evidence for spatial reorientation processes in human.

The difference in expected vs. unexpected starting alleys suggests that participants practiced expectation for a default starting alley. Importantly, the timing of the teleportation events was fully predictable and the participants were aware of the fact that the destination of the teleportation is random. Therefore, in theory, they could have (consciously or unconsciously) inhibited the processes that led to expectations regarding the destination of the teleportation. EEG evidence tells, however, that they nonetheless established prior expectations for the destination. Further, the found topographic differences are not bottom-up correlates of alley texture processing (see Fig. S4) but results of top-down spatial orientation processes based on the visual information conveyed by the alley textures.

One interesting question deriving from the current study is whether reorientation would be faster based on geometry than on textural cues. Ample evidence suggests that geometry serves as the most important cue for reorientation both in animals Keinath et al. (2017) and in children Spelke et al. (2010). Non-geometrical cues – such as texture – are also important, especially if geometrical cues are unreliable Pecchia & Vallortigara (2010) or if language is available to the actor Xu et al. (2017). In the current experiment, the geometry served no reliable cue for reorientation; however, the environment could be easily modified in a way where not textural but geometrical cues would help the reorientation. Interestingly, the latency of the found effects assumes that the complex visual information was evaluated rapidly, arguably in the same time scale as simpler visual features. This is in line with results from another study Howe (2017) showing that complex natural scenes are identified as rapidly as simple line directions.

We did not find difference in decision times in the current experiment, which is likely attributable to our experimental design and not to reorientation in general. The environment was very simple: since teleportation destinations were always either the North or South alleys, the unexpected alley meant only one alley and not three that the environment would have otherwise made possible. Therefore, the unexpected location was only “moderately unexpected”. Further studies may explore how further uncertainty regarding the unexpected position would affect decision times. Also, note that participants were not instructed to make quick choices but to try to find an underlying rule for the placement of the feedback object. Further studies may include more stringent reaction time tasks.

Although EEG provides a valuable window to the temporal dynamics of cortical processes, it lacks spatial resolution. Further studies using fMRI and EEG co-registration could explore the neural underpinnings of this reorientation process. Earlier studies showed that the retrosplenial cortex plays an important role in maintaining an allocentric frame of reference in an egocentric viewpoint-based task Park & Chun (2009); Lin et al. (2015). These results are further supported by single cell evidence that revealed cells in the retrosplenial complex process information in both egocentric and allocentric reference frames Alexander & Nitz (2015). Furthermore, retrosplenial activity is often detected over the parieto-occipital sites of the scalp EEG Lin et al. (2015).

Teleportation of the physical body of a living organism is a favorite topic of science fiction, however, virtual teleportation is not science fiction but an efficient way of locomotion in VR environments. The introduction of teleportation (or wormholes) means a violation to the Euclidean laws of geometry. Since the real world obeys Euclidean principles, one would easily assume that the human cognitive system does so as well. Interestingly though, not only people are able to learn environments with non-Euclidean geometries (e.g. wormholes Warren et al. (2017)) but they are often unaware of such violation. Moreover, a recent study by Vass et al. Vass et al. (2016) showed that when participants know the start and end points of such wormholes, they navigate them just like real routes as was apparent from the unattenuated theta oscillation in their hippocampi. These results suggest that the human brain maintains a flexible, graph like cognitive map Chrastil & Warren (2014); Tversky (1993) of the environment. To extend these findings, the current study showed EEG evidence for processing the mismatch between expected and unexpected destinations can be found as early as 100 msec.

Considering all these results, we can provide important practical directions for teleportation-based locomotion methods in virtual reality. It seems, that not only temporal but temporal-spatial predictability is an important prerequisite of effective teleportation. Otherwise, if teleportation is either spatially (see current results) or temporally unpredictable, a reorientation process has to occur Jezek et al. (2011). If used effectively though, teleportation could open up new ways of environment design, such where the layout does not obey the Euclidean principles yet is easily learned and memorized by people.

## Acknowledgements

The study was supported by the Research and Technology Innovation Fund (KTIA AIK 12-1-2013-0037).

## Supplementary Material

**Figure S1:**
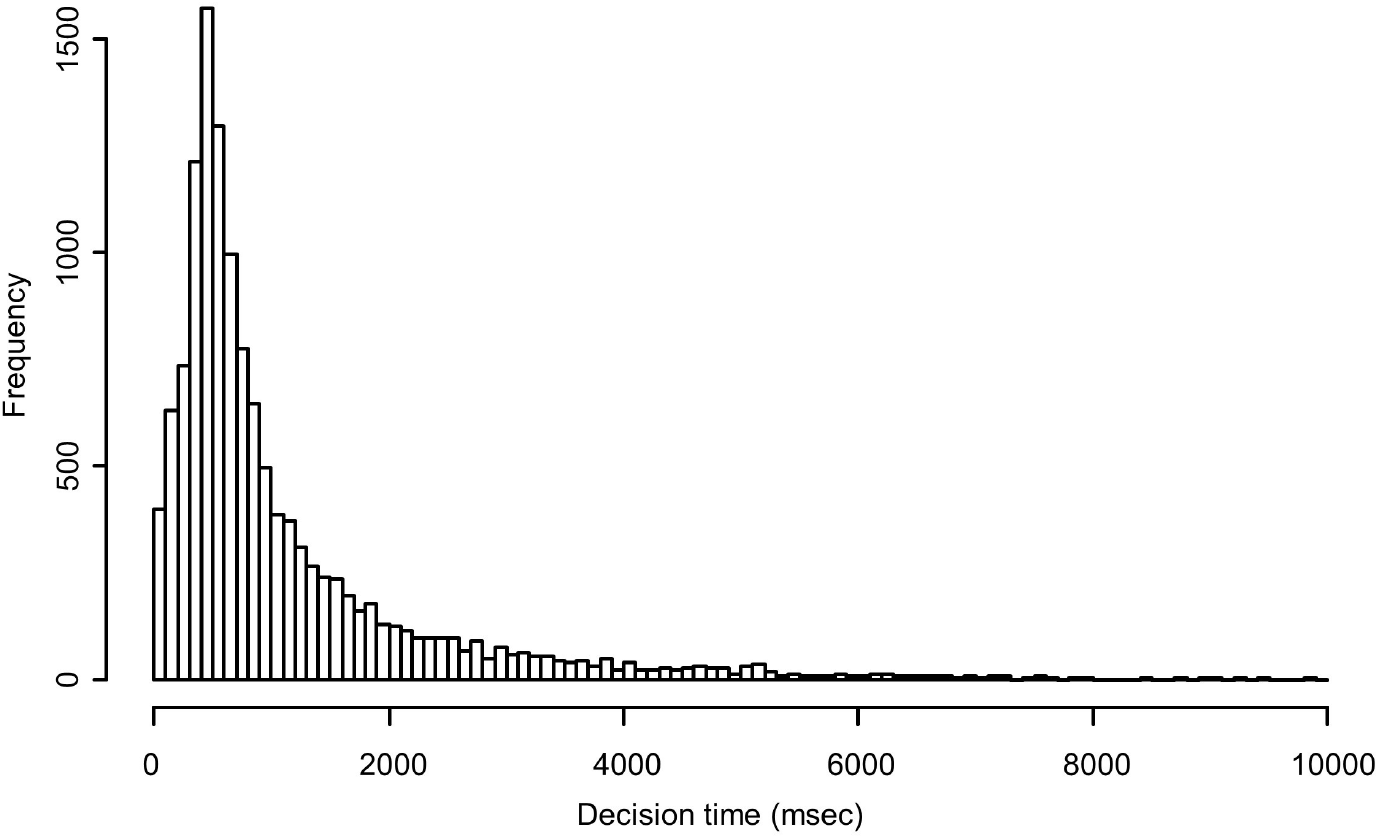
Distribution of decision times. As can be seen in the figure, the distribution of the decision times in the experiment showed a strongly left skewed distribution both when examined collapsed on all subjects (2.766) and when examined on the single subject level (*M_skewness_* = 2.801, *Min_skewness_* = 1.018, *Max_skewness_* = 9.923). Therefore, we calculate the median instead of the mean of the distributions. Nonetheless, the pattern of results remains when mean is calculated (*t* (33)=0.910, *p* = .815).

**Figure S2:**
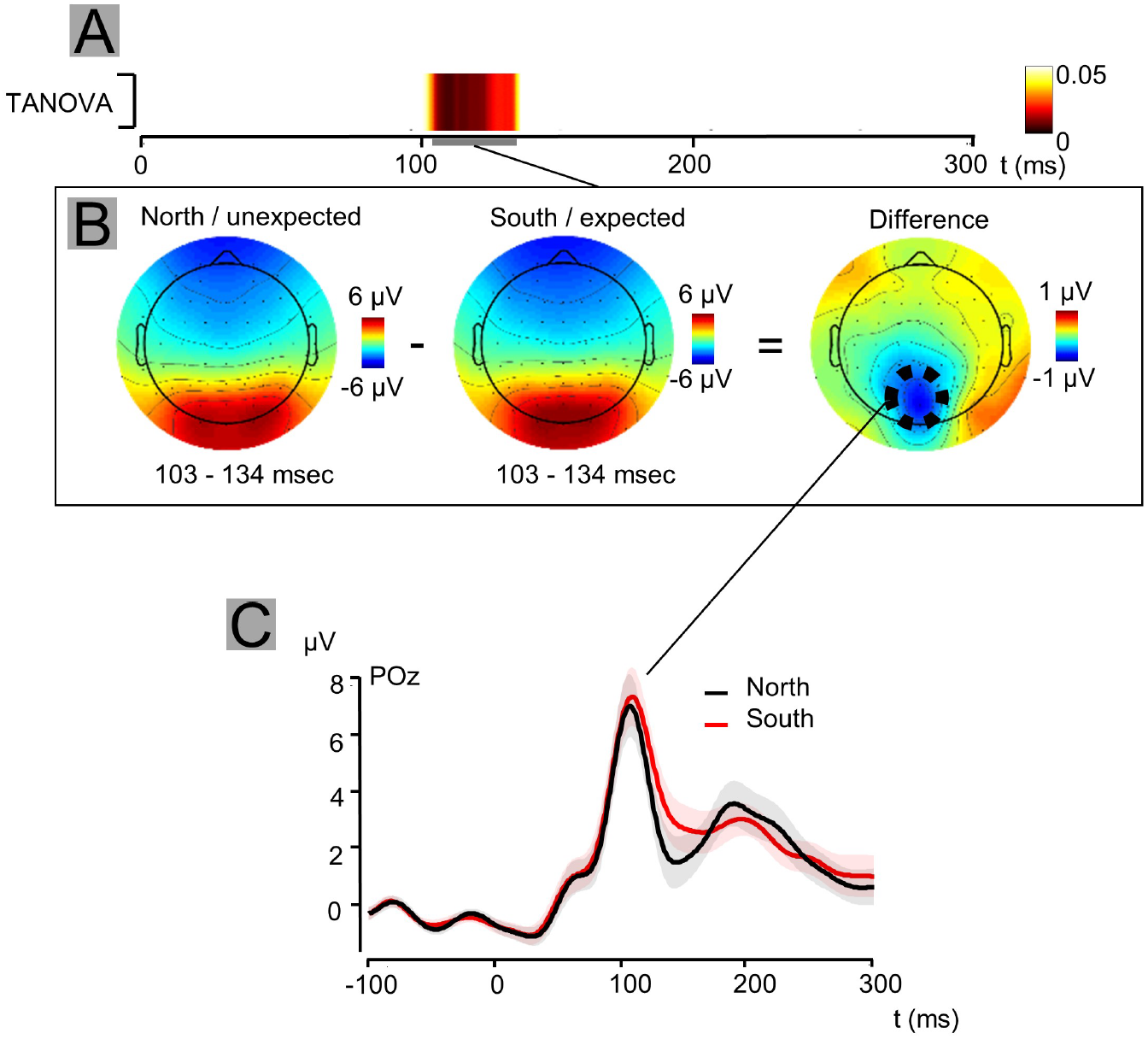
Analysis of the TANOVA maxima. A) TANOVA analysis revealed significant differences in topography between the unexpected and expected starting location at time points from 103 to 134 msec. B) The difference between topographies was a negative deflection over parieto-occipital sights, with a maximum on POz. C) Analysis on this electrode revealed significant difference between 114 and 163 msec and between 179 and 197 msec. These differences were significant both after FDR and cluster correction.

**Figure S3:**
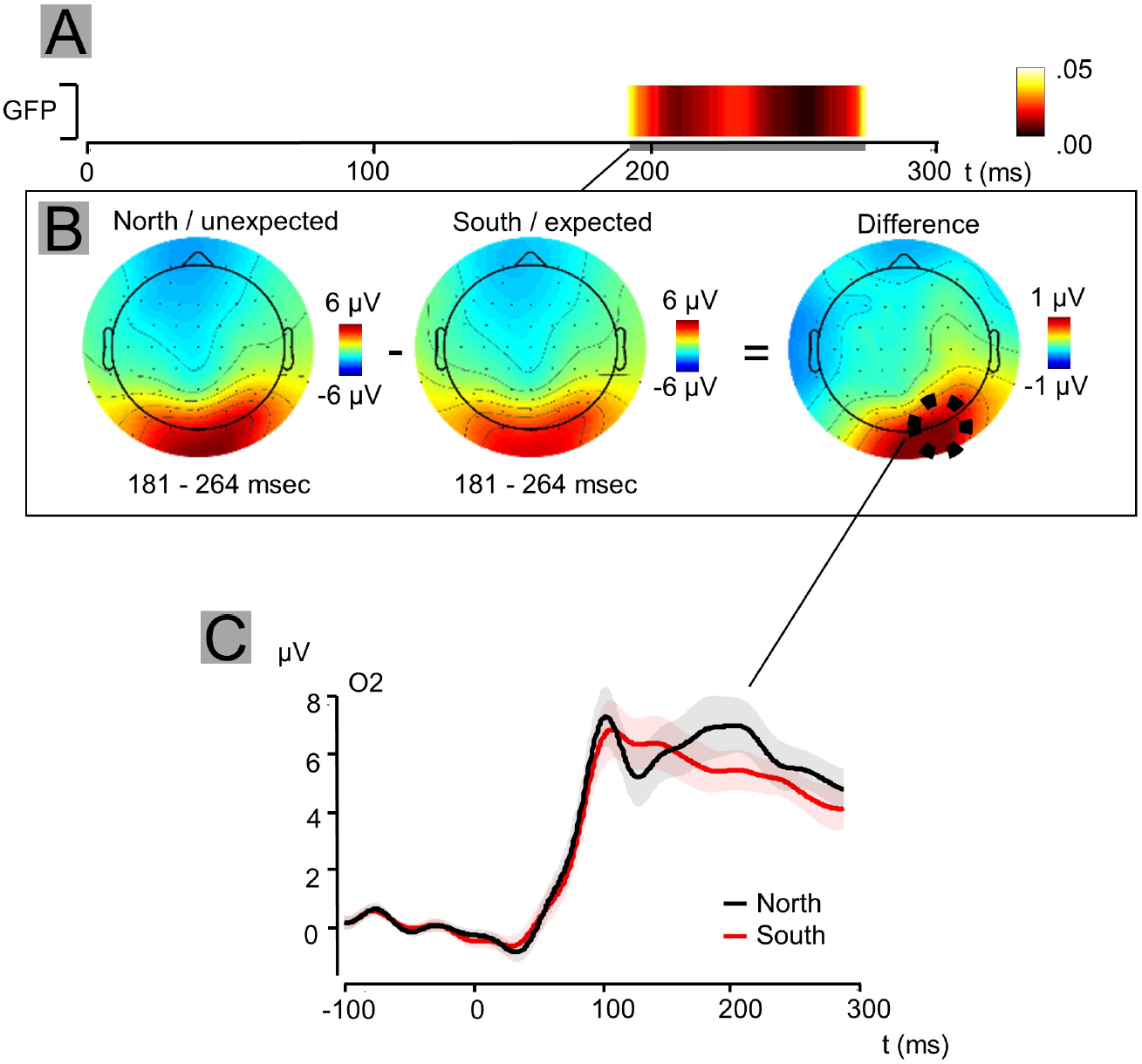
Analysis of the GFP maxima. A) GFP analysis revealed significant differences in topography between the unexpected and expected starting location at time points from 181 to 264 msec. B) The difference between topographies was a positive deflection slightly right from the occipital midline, with a maxima on O2. C) Analysis on this electrode revealed significant difference between 123 msec and 152 msec and between 175 and 300 msec. These differences were significant both after FDR and cluster correction.

**Figure S4:**
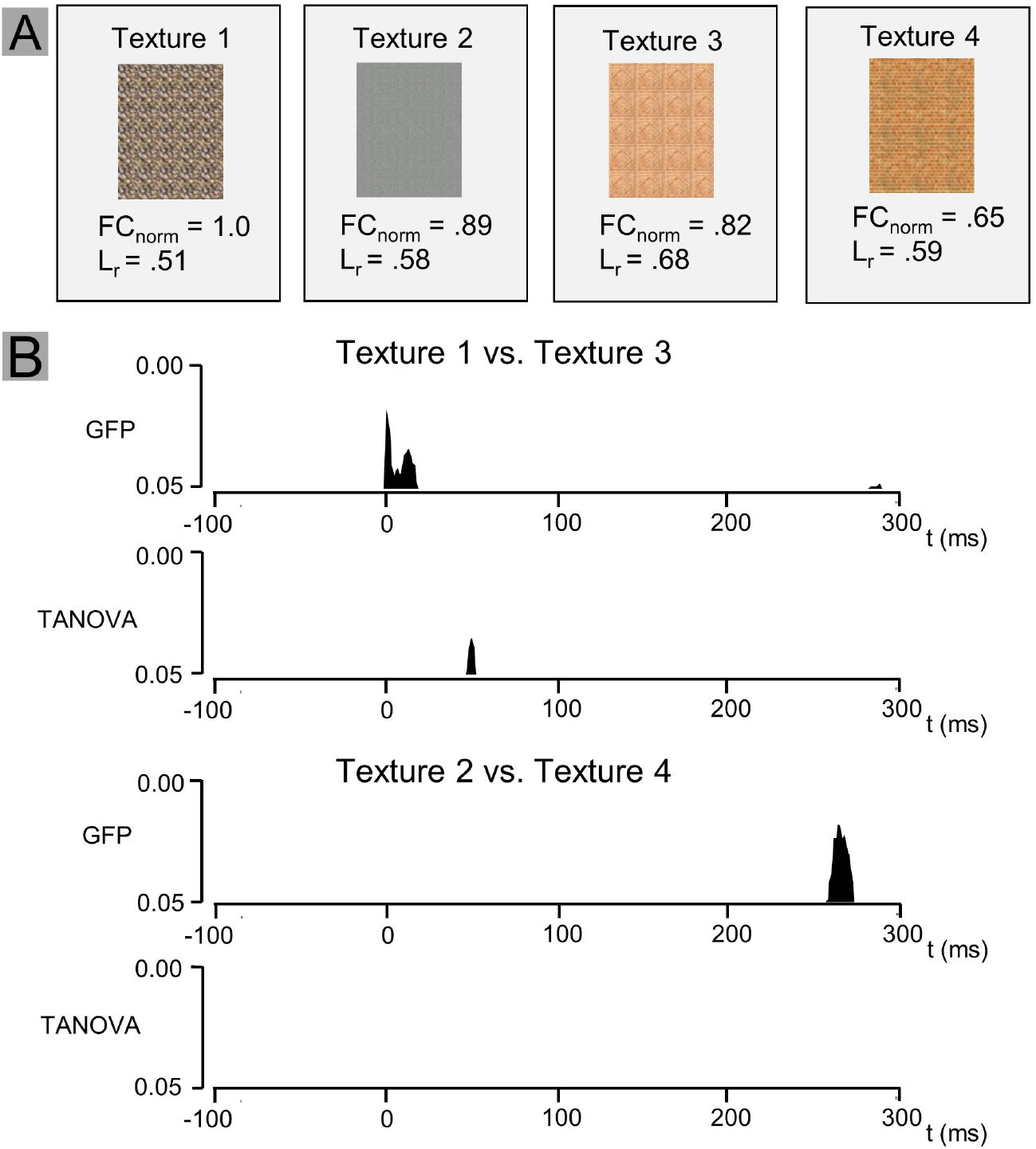
Control analysis of comparing the EEG to different textures in the starting alleys. the analysis of EEG data with the comparison of ERPs to the different destination alley textures. Since we randomly rotated the order of the wall textures for each participant and four textures were used, we were able to create two groups. In the first group, Texture 1 or Texture 3 were seen at the beginning of each trial, in the second group, Texture 2 and 4 were seen. We also quantified the visual complexity of the different textures by calculating their Feature Congestion Rosenholtz et al. (2007); based on this, the matched textures were similarly different in complexity for both groups. We ran separate analysis of topographic dissimilarity and global field power on both groups using the same parameters that we used during the main analysis. This analysis did not yield significant results in either group. Thus, we found no evidence for reorientation process that would have been driven by visual differences alone.

